# Predictions of the SARS-CoV-2 Omicron Variant (B.1.1.529) Spike Protein Receptor-Binding Domain Structure and Neutralizing Antibody Interactions

**DOI:** 10.1101/2021.12.03.471024

**Authors:** Colby T. Ford, Denis Jacob Machado, Daniel A. Janies

## Abstract

The genome of the SARS-CoV-2 Omicron variant (B.1.1.529) was released on November 22, 2021, which has caused a flurry of media attention due the large number of mutations it contains. These raw data have spurred questions around vaccine efficacy. Given that neither the structural information nor the experimentally-derived antibody interaction of this variant are available, we have turned to predictive computational methods to model the mutated structure of the spike protein’s receptor binding domain and posit potential changes to vaccine efficacy. In this study, we predict some structural changes in the receptor-binding domain that may reduce antibody interaction without completely evading existing neutralizing antibodies (and therefore current vaccines).

## Introduction

A team of researchers from the Botswana-Harvard HIV Reference Laboratory submitted a new SARS-CoV-2 genome sequence to GISAID on November 22, 2021 (GISAID accession. EPI_ISL_6752027). The specimen was taken from a living 59-year-old male from Gaborone, Botswana using a nasopharyngeal swab and was sequenced using a Nanopore MinION device.

This sample’s genome contains 60 mutations from the Wuhan-derived reference genome (GenBank accession no. NC_045512.2) (1), 37 of which are in the Spike (S) protein. This variant was given the identifier B.1.1.529 by PANGO lineages (2). On November 26, 2021, the WHO has designated B.1.1.529 as a Variant of Concern (VOC), named Omicron (3).

The emergence of new SARS-CoV-2 variants is expected. Therefore, scientists have advocated for close international monitoring to determine the need for vaccination boosters and redesign (4). Hence, the identification of the omicron variant is not surprising. What is surprising is the number of mutations that the omicron variant accumulated compared to the first sequenced genome of SARS-CoV-2.

Different authors have warned that limited SARS-CoV-2 sampling and sequencing from positive cases, especially from asymptomatic and symptomatic cases that did not require hospitalization, would make it challenging to identify new mutations in the virus. For example, Brito et al. (5) analyzed the spatiotemporal heterogeneity in each country’s

SARS-CoV-2 genomic surveillance efforts using metadata submitted to GISAID until May 30, 2021. These authors calculate that sequencing capacity should be at least 0.5% of cases per week when incidence is more than 100 positive cases every 100,000 people. Unfortunately, most countries are not reaching this sequence threshold.

While sampling bias can explain why we may miss new mutations and fail to identify new variants of low prevalence, the emergence of new variants is due to factors that favor the transmission of SARS-CoV-2, including low vaccination rates in some regions, especially in low and middle-income countries (LMICs). Therefore, disparities in vaccination rates combined with sampling bias explain why scientists may continue to be surprised by the mutations in new SARS-CoV-2 variants.

There are many questions regarding genomic epidemiology and the lessons we can learn from the COVID-19 pandemic. Those questions are beyond the scope of this manuscript, and we addressed them elsewhere (6). While the origin and evolution of the Omicron variant are still open questions, here we focus on the potential implications of the mutations observed in this variant.

This seemingly hyper-mutated variant is of public health concern with unanswered questions surrounding vaccine protection (from available vaccines), the possibility of reinfection, transmissibility, and pathogenicity.

Regarding vaccine efficacy, we must look at the receptor-binding domain (RBD), part of the S1 subunit, of the spike protein as this is the binding site for neutralizing antibodies. This domain exists between positions 319 and 541 of the spike protein. Omicron contains 15 mutations in the RBD, none of which are deletions or insertions. In contrast, the Delta variant contains 7 mutations across the entire spike protein, only 2 of which are in the RBD.

Given that an experimentally-derived structure of the Omicron spike protein is not yet available, we must derive a predicted structure from its sequence *in silico*. Then, we can use available neutralizing antibody structures to computationally model the interaction between Omicron and the paratopes of the antibodies, thus allowing us to compare potential affinity changes due to the mutations and posit their effects to vaccine efficacy.

## Methods

### Sequence Comparison among VBMs and VOCs

We downloaded the reference genome of SARS-CoV-2 (WuhanHu-1, NCBI’s RefSeq accession no. NC_045512.2) as well as the first 100 complete genome sequences (≥ 29,000 bp) of each Variant of Concern (VOC) and Variant Being Monitored (VBM). The total number of input sequences was 1,301. We aligned all of these complete genomes using `MAFFT` version 7.475 (7) with the “auto” option and trimmed the alignment to remove the 5’-UTR and 3’-UTR regions. We also removed duplicated sequences or sequences with more than 5% of missing data, leaving us with 1,026 sequences.

We annotated each of the 1,026 remaining sequences using the strategy described in Machado et al. (2021) (8). Once we had all the predicted spike proteins for each of the 1,026 genomes, we aligned those sequences based on their translation with the help of `MAFFT` using the `TranslatorX` pipeline (9). We removed duplicated sequences and sequences with more than 5% of missing data. Finally, we identified the receptor binding motif on that alignment based on sequence similarity with the reference.

We then calculated the pairwise p-distances between each pair of sequences were calculated using `MEGA` version 11.0.10 (10). This distance is the proportion (p) of nucleotide sites at which two sequences being compared are different. The p-distances were calculated for the whole spike alignments (nucleotides) but also for the alignment of its receptor binding motif (RBM; position 430–522 of the spike amino acid sequence, a subset of the positions in the RBD).

This variant nucleotide sequence for the spike protein was then translated into amino acids using the standard translation table. This sequence was then trimmed to only contain the RBD of the spike protein (positions 319 to 541).

### Receptor-Binding Domain Structural Prediction

Using the derived RBD amino acid sequence for Omicron, we used `AlphaFold2` and `RoseTTAFold` to create a predicted 3D protein structures. `AlphaFold2` is a neural network-based deep learning model created by Google DeepMind (11). The algorithm first searches for homologous sequences with existing structures to use as a scaffold on which to place the new sequence. `RoseTTAFold` is a similar neural network-based system by the Institute for Protein Design at the University of Washington (12).

The `AlphaFold2`-based prediction was run with the “single sequence” mode using the predicted TM-score (PTM) method. We also specified that the algorithm should run an Amber relaxation procedure to repair any structural violations in the predicted model (13). The `RoseTTAFold`-based prediction was run with the “mmseqs2” mode (by ColabFold (14)).

Both systems each resulted in a .PDB file of the predicted RBD structure for Omicron along with metrics surrounding the multiple sequence alignment coverage, predicted aligned error (PAE), and predicted confidence (pLDDT) by position (available in Supplementary Materials).

Given that this study focuses on antibodies that bind to the top of the RBD of the spike protein. Since that `AlphaFold2` and `RoseTTAFold` are template-based models that generate the predicted RBD structures using homologous sequences for which we have actual structures, we can avoid modeling the entire spike protein.

### Neutralizing Antibody Interaction Simulation

Using the predicted structures of the Omicron RBD, we simulated the interaction with four available neutralizing antibody structures: C105, CC12.1, CC12.3, and CV30 (PDBs: 6XCM, 6XC2, 6XC7, and 6XE1, respectively) (15–17). We used only a single fragment antigen-binding (Fab) region of the antibody structures as the paratope location against which to dock. Each of the RBD structures from these reference files have identical sequences to the Wuhan-Hu-1 spike RBD. Each of these neutralizing antibody structures were collected from patients who had been infected with SARS-CoV-2. Thus, these structures are serologically-derived antibodies rather than structures of therapeutic antibodies. All of them bind to the same “up” location of the S1 subunit of the spike protein (class I binders). This is a similar location to the interaction site between the human ACE2 receptor epitope. Thus, the neutralizing mechanism of these antibodies is in the prevention of SARS-CoV-2 binding to ACE2 on human cells.

We used `HADDOCK` version 2.4, a biomolecular modeling software that provides docking predictions for provided structures, to predict the binding affinity between the epitope of the RBD with the paratope of the neutralizing antibody structures (18). This takes in two or more .PDB files as inputs and outputs multiple predicted protein complexes in .PDB format along with docking metrics.

We first renumbered the residues according to `HADDOCK`’s requirements and then specified the interacting residues between the RBD structure and the Fab. Specifically, ensure there are not overlapping residue IDs between the chains of a .PDB file and then specify the residues that are assumed to interact between the structures. This analysis was performed on the antibody-RBD structure pairs shown in Table 1.

**Table 1.** List of analyses performed, comparing reference and predicted RBD structures in complex with reference Fab structures.

| Antibody Fab | Analysis Type | RBD Source |
| --- | --- | --- |
| C105<br>(6XCM, chains N and S) | Reference | 6XCM<br>(chain B) |
|  | Prediction | B.1.1.529<br>(from AlphaFold2) |
|  | Prediction | B.1.1.529<br>(from RoseTTAFold) |
| CC12.1<br>(6XC2, chains H and L) | Reference | 6XC2<br>(chain A) |
|  | Prediction | B.1.1.529<br>(from AlphaFold2) |
|  | Prediction | B.1.1.529<br>(from RoseTTAFold) |
| CC12.3<br>(6XC7, chains C and D) | Reference | 6XC7<br>(chain A) |
|  | Prediction | B.1.1.529<br>(from AlphaFold2) |
|  | Prediction | B.1.1.529<br>(from RoseTTAFold) |
| CV30<br>(6XE1, chains H and L) | Reference | 6XE1<br>(chain B) |
|  | Prediction | B.1.1.529<br>(from AlphaFold2) |
|  | Prediction | B.1.1.529<br>(from RoseTTAFold) |

The assessment of these interactions was measured by multiple biophysical factors including van der Waals energy, electrostatic energy, desolvation energy, and restraints violation energy, which were collectively used to derive a HAD-DOCK score to quantify changes in protein-protein interaction resulting from mutations in the RBD. Further, interfacing residues between the RBD and Fab structures were determined by identifying residues above the 1.0 Å^2^ difference in surface area cutoff between the chains of the RBD and the Fab using the *InterafaceResidues*^1^ functionality in `PyMol` version 2.4.1 (19).

We then compared the metrics of the actual complexes (i.e., the real RBD structure and the Fab) versus the predicted RBD structures of Omicron (with the same Fab). This provides a baseline interaction that was then measured against the mutated interactions with each respective Fab.

Differences between HADDOCK results were assessed using Kruskal-Wallis tests and ad hoc Wilcoxon pairwise comparisons. All statistical analyses were performed using `R` version 4.0.4 (20).

## Results

### Variant Sequence Comparison

Although the Omicron RBM (spike amino acid sequence, positions 430–522) can be efficiently categorized by nine characteristic mutations (S:N440K, S:G446S, S:S447N, S:T478K, S:E484A, S:Q493R, S:G496S, S:Q298R, S:N501Y), at least two of them (S:N440K and S:G446S) may be missing from some samples classified as Omicron. Also, some Omicron RBMs contains an additional mutation at S:Y505H.

The Omicron variant is the variant more distantly related to the reference genome (SARS-CoV-2 Wuhan-Hu-1; NCBI’s RefSeq accession no. NC_045512.2) in the proportion of shared nucleotides. Also, Omicron is the variant that is more distantly related to Gamma. See Figures 2 and 3.

**Fig. 1.**
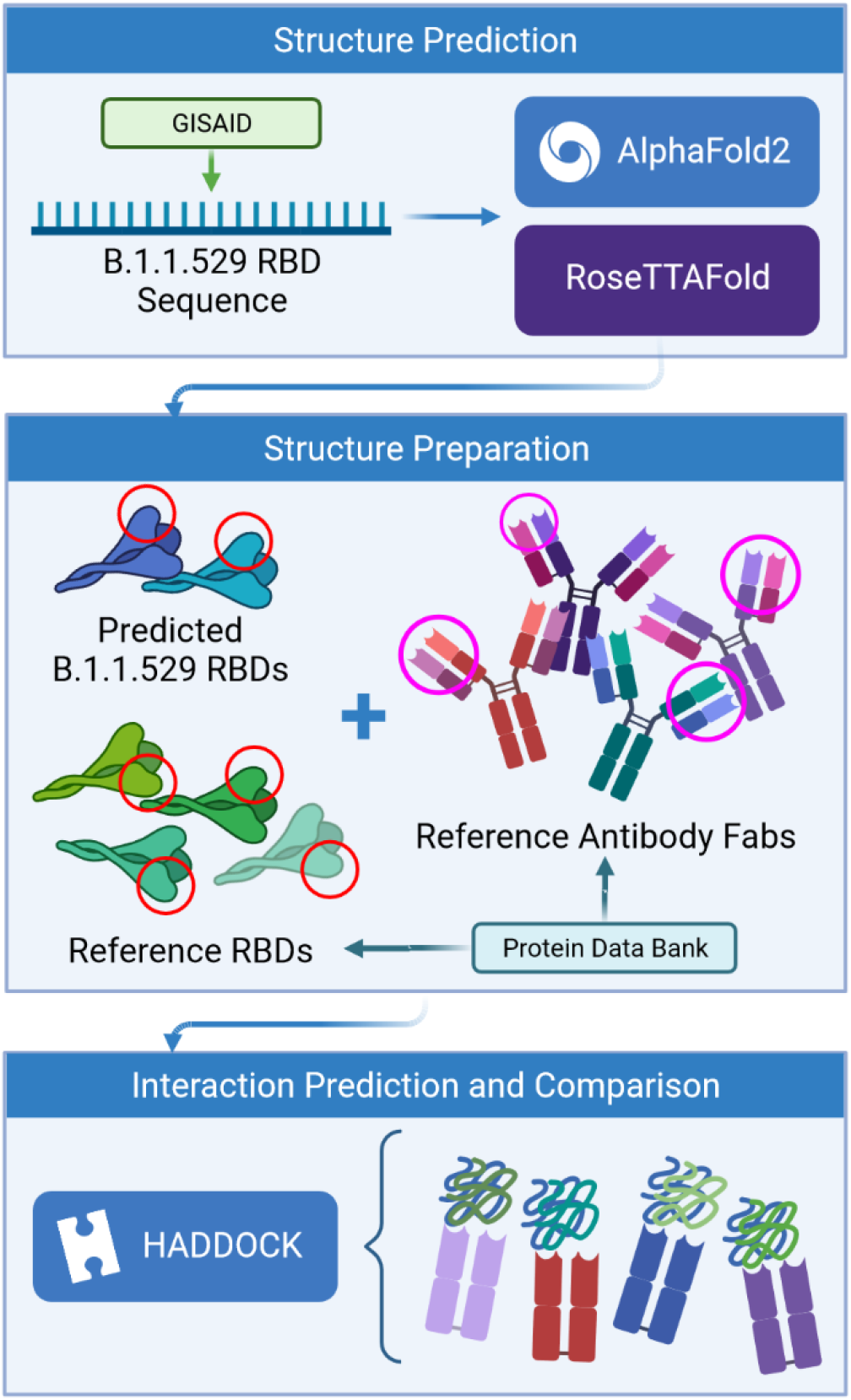
Process flow of the prediction analysis steps.

**Fig. 2.**
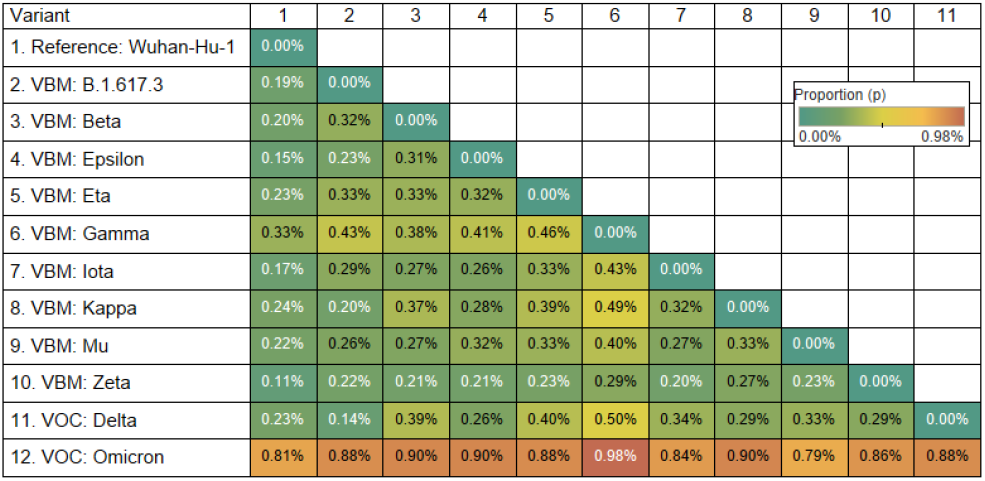
Distance matrix of the spike gene (using nucleotides) for 9 Variants Being Monitored (VBM) and 2 Variants of Concern (VOC). The distance is the average proportion (*p*) of nucleotide sites at which two sequences being compared are different.

**Fig. 3.**
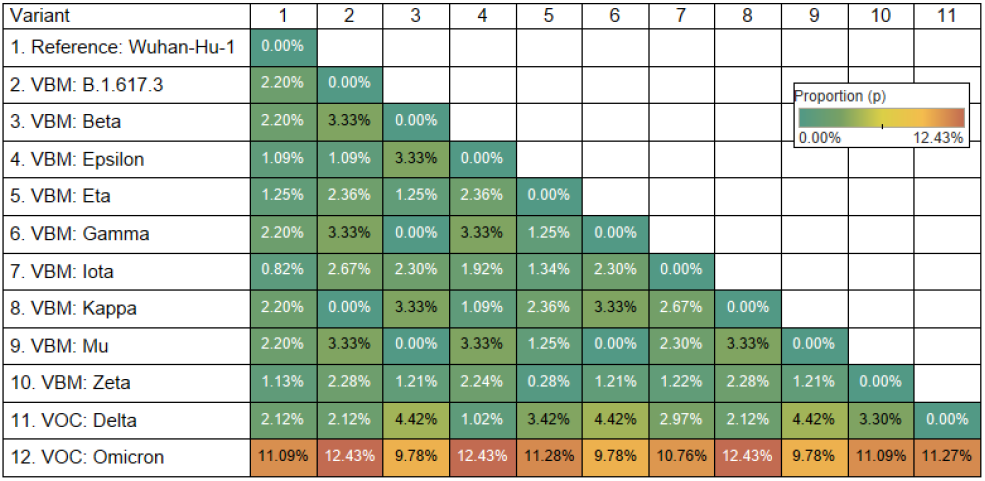
Distance matrix of the receptor binding motif (RBM) of spike gene (using amino acids) for 9 Variants Being Monitored (VBM) and 2 Variants of Concern (VOC). The distance is the average proportion (*p*) of nucleotide sites at which two sequences being compared are different.

### Mutational Analysis

Comparing the RBD of Omicron to the reference genome, there are 15 mutations, all of which are single amino acid substitutions. Most of the substitutions result in a change in the residue type. (See Table 2.)

**Table 2.**
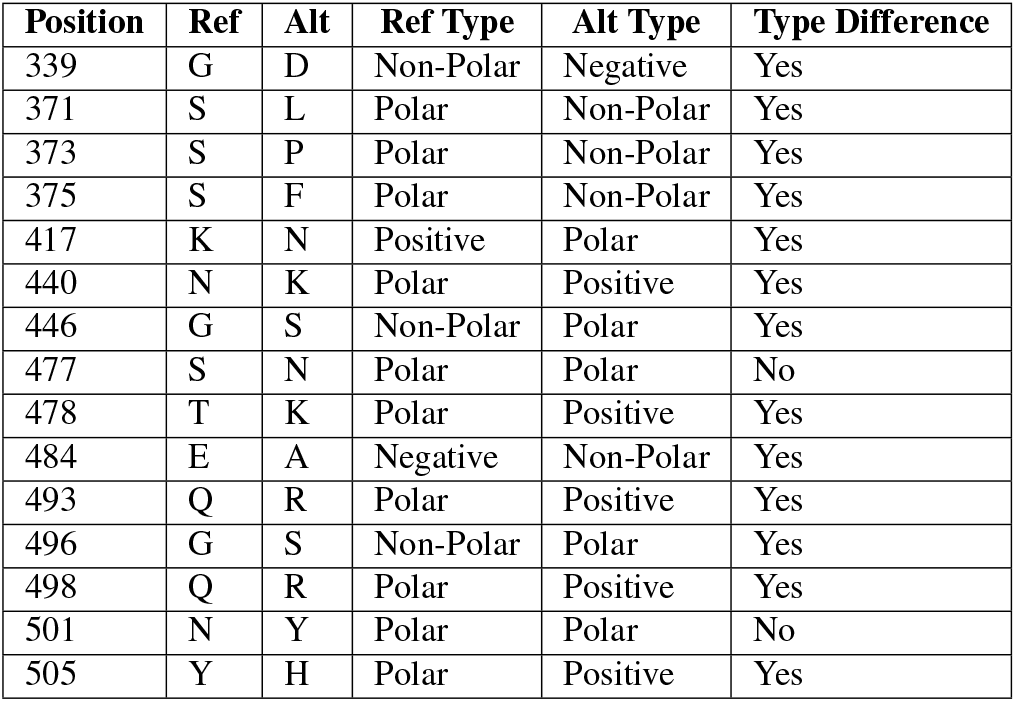
Mutations in the receptor-binding domain (RBD) of the spike protein in the Omicron variant (B.1.1.529).

The resulting RBD structure from `AlphaFold2` predicts that there is little conformational change from the reference structure. Conversely, there is a significant conformational change in the predicted RBD structure from `RoseTTAFold`. See Figure 4.

**Fig. 4.**
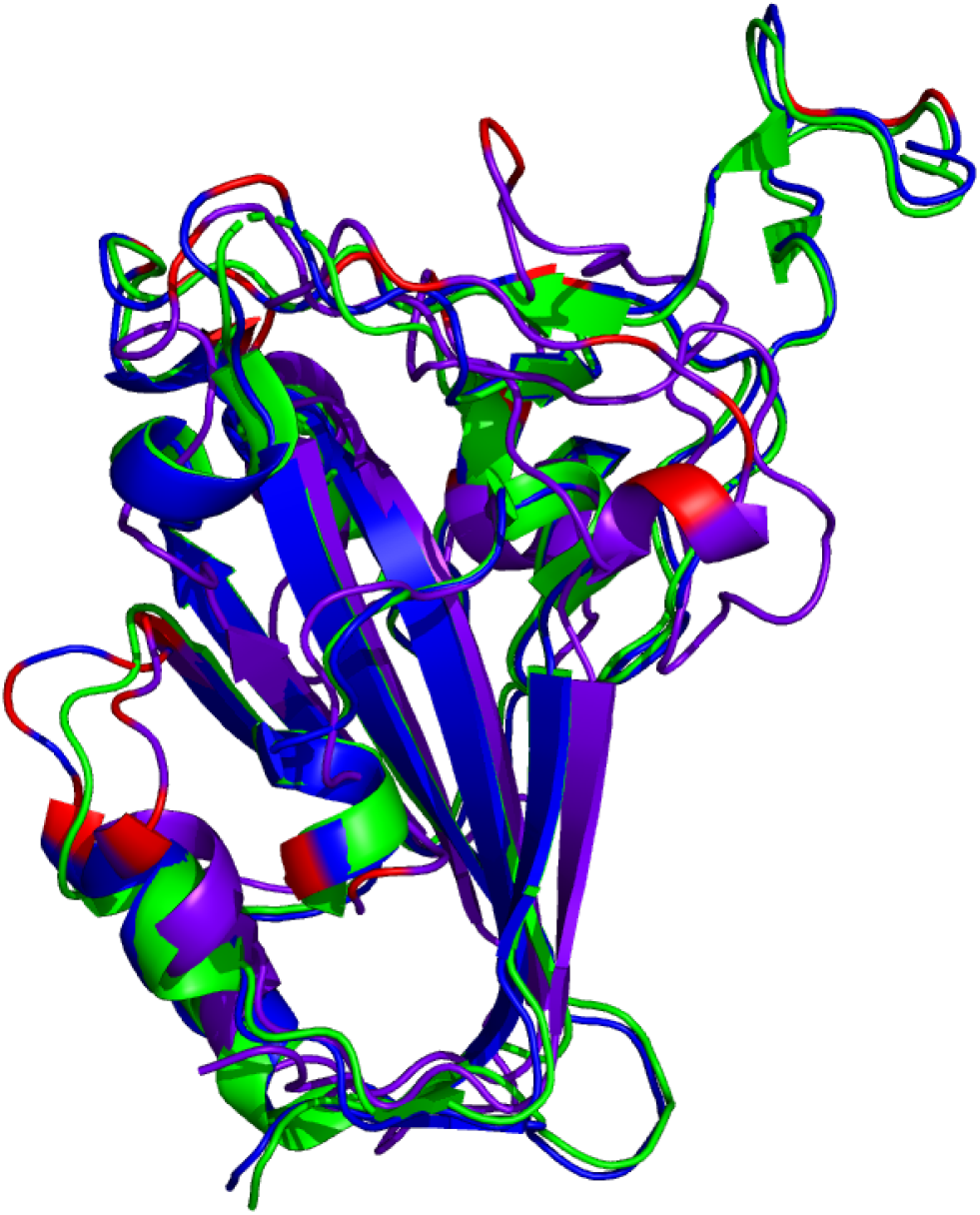
Comparison of reference RBD structure (PDB: 6XC2, shown in green) and the predicted Omicron (B.1.1.529) RBD structures (`AlphaFold2` shown in blue, `RoseTTAFold` shown in purple). Mutated residues are highlighted in red.

There are multiple mutated residues (shown in red in Figures 5 and 6) in positions that may affect the ability of a neutralizing antibody to sufficiently bind. Some of these mutated residues change to much longer side-chained or differently-charged amino acids. For example, there are two “to lysine” mutations: N440K and T478K (i.e., from polar, smaller side chain residues to a positive-charged, longer side chain residue). These types of changes may have an effect on the binding affinity between the RBD and an antibody, either by changing the surface charge on the protein or by inhibiting a tighter antibody interaction.

**Fig. 5.**
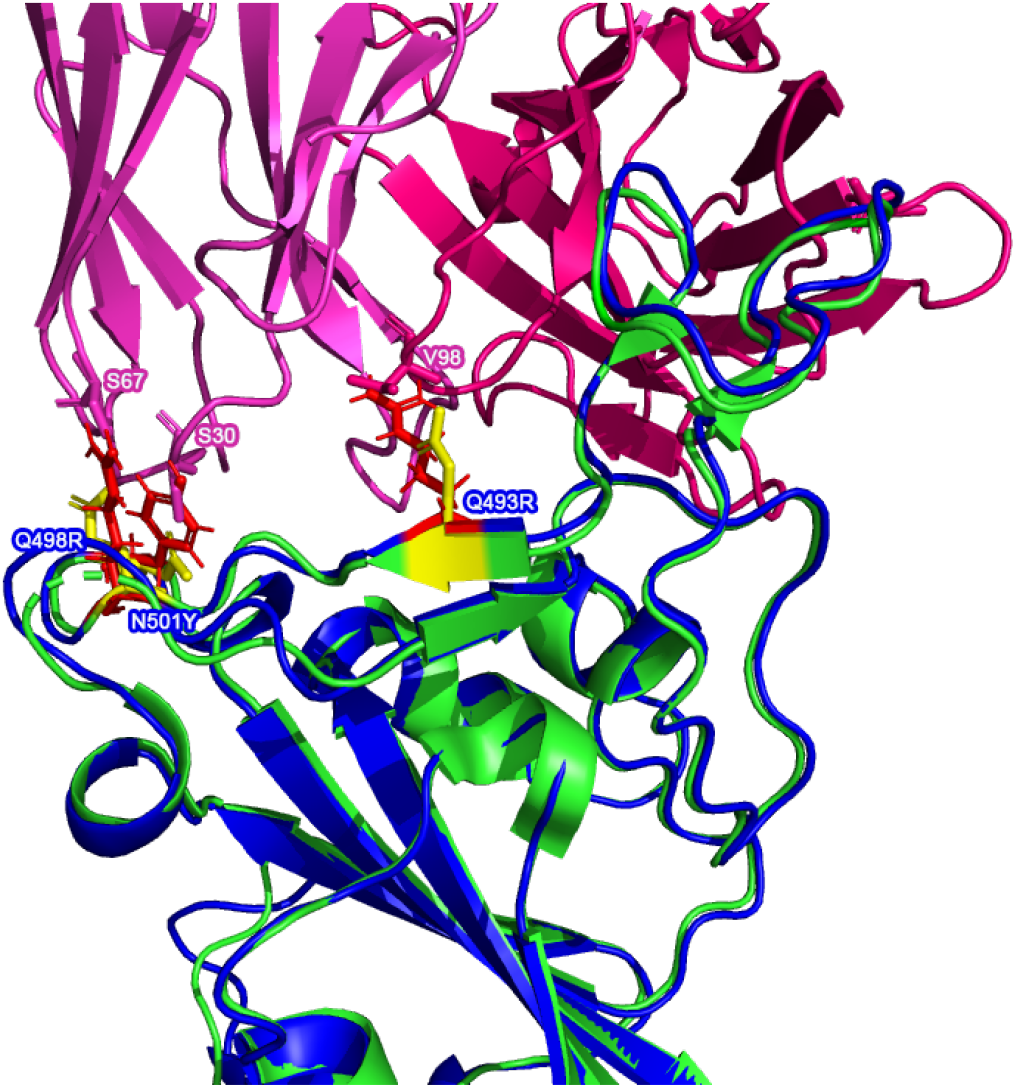
Possible inhibitory mutated RBD residues in the `AlphaFold2` Omicron (B.1.1.529) (structure shown in blue with mutated residues of interest shown in red) superimposed on a reference RBD structure. Note: the reference RBD structure (PDB: 6XC2) is shown in green with equivalent position residues highlighted in yellow. CC12.1 antibody Fab (from PDB 6XC2) is shown in magenta / pink.

**Fig. 6.**
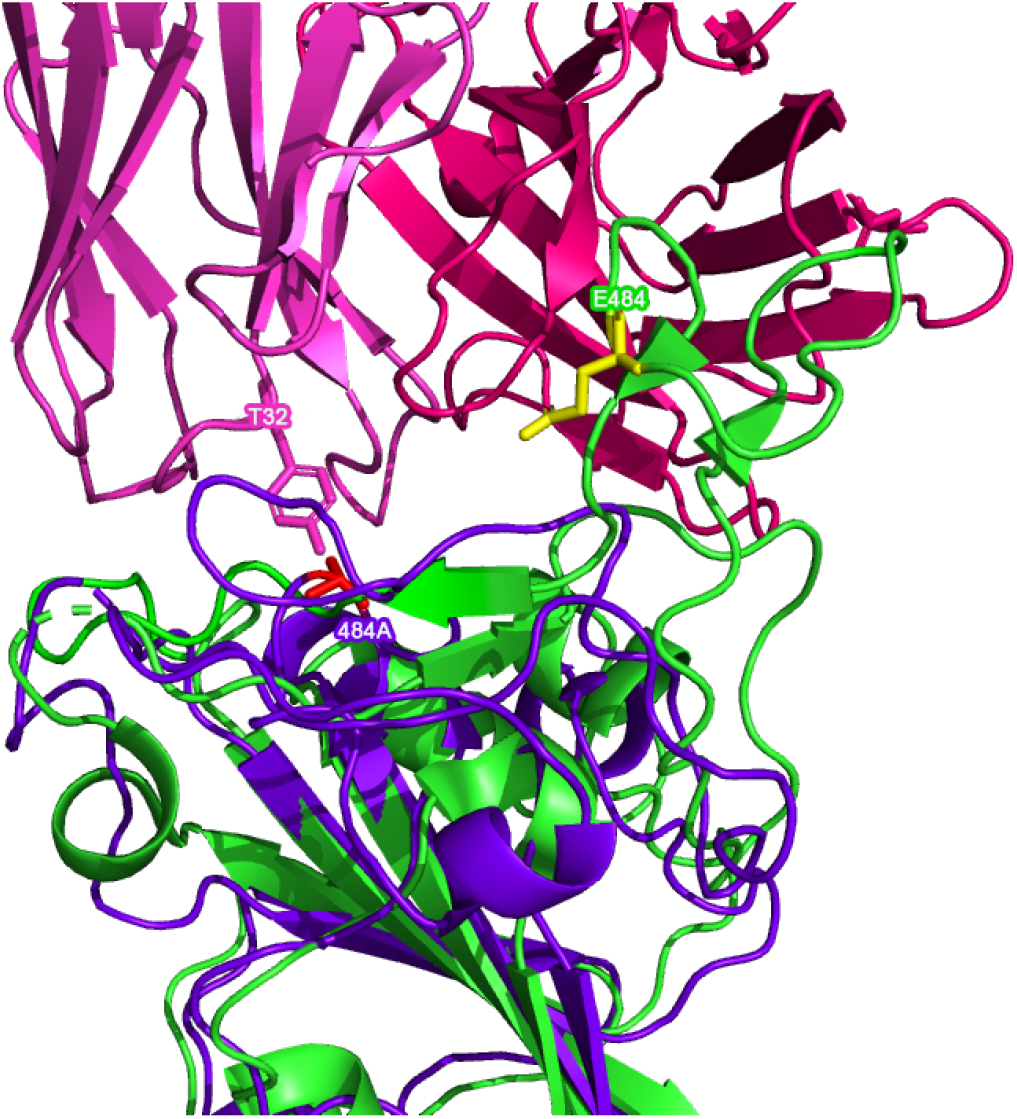
Possible inhibitory mutated RBD residues in the `RoseTTAFold` Omicron (B.1.1.529) (structure shown in purple with mutated residues of interest shown in red) superimposed on a reference RBD structure. Note: the reference RBD structure (PDB: 6XC2) is shown in green with equivalent position residues highlighted in yellow. CC12.1 antibody Fab (from PDB 6XC2) is shown in magenta / pink.

Superimposing the predicted RBDs on the reference RBD with the CC12.1 antibody in place, shown in Figures 5 and 6, shows that mutations Q493R, Q498R, and N501Y in the `AlphaFold2` structure and the E484A mutation in the `RoseTTAFold` structure may affect the binding position of the antibody. These side chains clash with particular residues of the Fab, which may cause a less effective and more distant binding. Longer/larger side chains may increase the distance between the Fab paratope of the antibody and the epitope of the RBD.

### Antibody Binding Analyses

The results of all four antibody docking exercises with the predicted `AlphaFold2` RBD structure show that the Fab of the respective neutralizing antibodies continue to bind to the RBD of Omicron, though not as well as the reference interaction. Note that there is a consistent decrease (increase in value) in the electrostatic energy and an increase in restraints violation energy between the binding from the reference RBDs and the predicted RBD of Omicron. The HADDOCK score is worse (higher) across the board and it appears that the interaction of the Omicron RBD with the antibodies are more distant, as shown by the buried surface area changes below.

When looking at the antibody docking exercises with the predicted `RoseTTAFold` RBD structure, there is agreement with the `AlphaFold2` results in that there still seems to be interaction with the neutralizing antibodies. However, the reduction in binding affinity is much more severe given the conformational changes seen only in the `RoseTTAFold` structure.

### Note

All values in Tables 3, 4, 5, and 6 below represent the best docking predictions from `HADDOCK`. Also, values in parentheses represent the percentage difference between the given metric for the predicted structure and the reference structure.

**Table 3.** HADDOCK metrics for the C105 docking prediction, comparing the 6XCM RBD vs. the Omicron (B.1.1.529) predicted RBD structures.

| Metric | 6XCM RBD w/ C105 | Predicted B.1.1.529 RBD w/ C105 |  |
| --- | --- | --- | --- |
|  |  | AlphaFold | RoseTTAFold |
| van der Waals energy | -85 | -71.8 (16%) | -44.2 (48%) |
| Electrostatic energy | -280.9 | -268.9 (4%) | -324.5 (-16%) |
| Desolvation energy | -18.6 | -17.1 (8%) | -3.2 (83%) |
| Restraints violation energy | 154.1 | 158.8 (3%) | 338.6 (120%) |
| HADDOCK score | -144.4 | -126.8 (12%) | -78.4 (46%) |
| Buried Surface Area | 2417.8 | 2299.6 (-5%) | 1555.1 (-36%) |

**Table 4.** HADDOCK metrics for the CC12.1 docking prediction, comparing the 6XC2 RBD vs. the Omicron (B.1.1.529) predicted RBD structures.

| Metric | 6XC2 RBD w/ CC12.1 | Predicted B.1.1.529 RBD w/ CC12.1 |  |
| --- | --- | --- | --- |
|  |  | AlphaFold | RoseTTAFold |
| van der Waals energy | -106.9 | -90.7 (15%) | -27.5 (74%) |
| Electrostatic energy | -342.6 | -284.9 (17%) | -259.2 (24%) |
| Desolvation energy | -37.5 | -28.8 (23%) | -12.4 (67%) |
| Restraints violation energy | 143.2 | 152.6 (7%) | 201.4 (41%) |
| HADDOCK score | -198.6 | -163.3 (18%) | -71.6 (64%) |
| Buried Surface Area | 2778.5 | 2584.3 (-7%) | 1578.4 (-43%) |

**Table 5.** HADDOCK metrics for the CC12.3 docking prediction, comparing the 6XC7 RBD vs. the Omicron (B.1.1.529) predicted RBD structures.

| Metric | 6XC7 RBD w/ CC12.3 | Predicted B.1.1.529 RBD w/ CC12.3 |  |
| --- | --- | --- | --- |
|  |  | AlphaFold | RoseTTAFold |
| van der Waals energy | -86.1 | -93.1 (-8%) | -48.1 (44%) |
| Electrostatic energy | -248.1 | -194.1 (22%) | -189.2 (24%) |
| Desolvation energy | -41.7 | -37.9 (9%) | -23.5 (44%) |
| Restraints violation energy | 159.1 | 104.7 (-34%) | 221.4 (39%) |
| HADDOCK score | -161.5 | -159.4 (1%) | -87.4 (46%) |
| Buried Surface Area | 2489.8 | 2416.7 (-3%) | 1420.6 (-43%) |

**Table 6.**
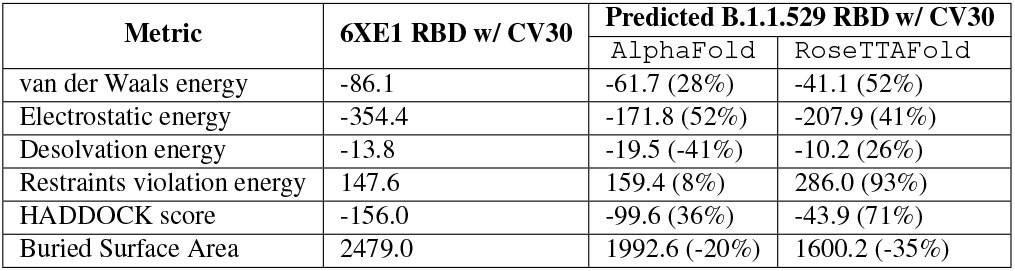
HADDOCK metrics for the CV30 docking prediction, comparing the 6XE1 RBD vs. the Omicron (B.1.1.529) predicted RBD structures.

#### C105 Antibody Binding

Resulting binding metrics from the C105 `HADDOCK` docking analysis are shown in Table 3. This interaction shows that there is a ∼; 4% reduction in the electrostatic energy and an ∼ 5% decrease in buried surface area comparing between the 6XCM RBD and the predicted RBD of Omicron from `AlphaFold2`. From `RoseTTAFold`, the interaction shows that there is ∼ 16% increase in the electrostatic energy but with a ∼ 36% decrease in buried surface area.

#### CC12.1 Antibody Binding

Resulting binding metrics from the CC12.1 `HADDOCK` docking analysis are shown in Table 4. This interaction shows that there is a ∼ 17% reduction in the electrostatic energy and an ∼ 7% decrease in buried surface area comparing between the 6XC2 RBD and predicted RBD of Omicron from `AlphaFold2`. From `RoseTTAFold`, the interaction shows that there is ∼ 24% reduction in the electrostatic energy but and a ∼ 43% decrease in buried surface area.

#### CC12.3 Antibody Binding

Resulting binding metrics from the CC12.3 `HADDOCK` docking analysis are shown in Table 4. This interaction shows that there is a ∼ 22% reduction in the electrostatic energy and an ∼ 3% decrease in buried surface area comparing between the 6XC7 RBD and predicted RBD of Omicron from `AlphaFold2`. From `RoseTTAFold`, the interaction shows that there is ∼ 24% reduction in the electrostatic energy and a ∼ 43% decrease in buried surface area.

#### CV30 Antibody Binding

Resulting binding metrics from the CV30 `HADDOCK` docking analysis are shown in Table 6. This interaction shows that there is a ∼ 52% reduction in the electrostatic energy and an ∼ 20% decrease in buried surface area comparing between the 6XE1 RBD and predicted RBD of Omicron from `AlphaFold2`. From `RoseTTAFold`, the interaction shows that there is ∼ 41% reduction in the electrostatic energy and a ∼ 35% decrease in buried surface area.

### Antibody Interaction Comparison

All of the interaction predictions among the four antibodies tested in this study (C105, CC12.1, CC12.3, and CV30) agree that there is a decrease in binding affinity when comparing the respective RBD interactions with the Omicron RBD interactions. Across all of the docking predictions using the `AlphaFold2` RBD structure, we see a drop in electrostatic interaction (increase in the electrostatic energy value) rang-ing from ∼ 4% to ∼ 52% and a consistent decrease in buried surface area (increase in distance) of the RBD and the antibody Fab. In addition, we see a variable increase (worsening) in the HADDOCK score, indicating that all of the Omicron RBD structures have a lower binding affinity when compared to their respective reference RBD structures as a benchmark. See Figure 11.

**Fig. 7.**
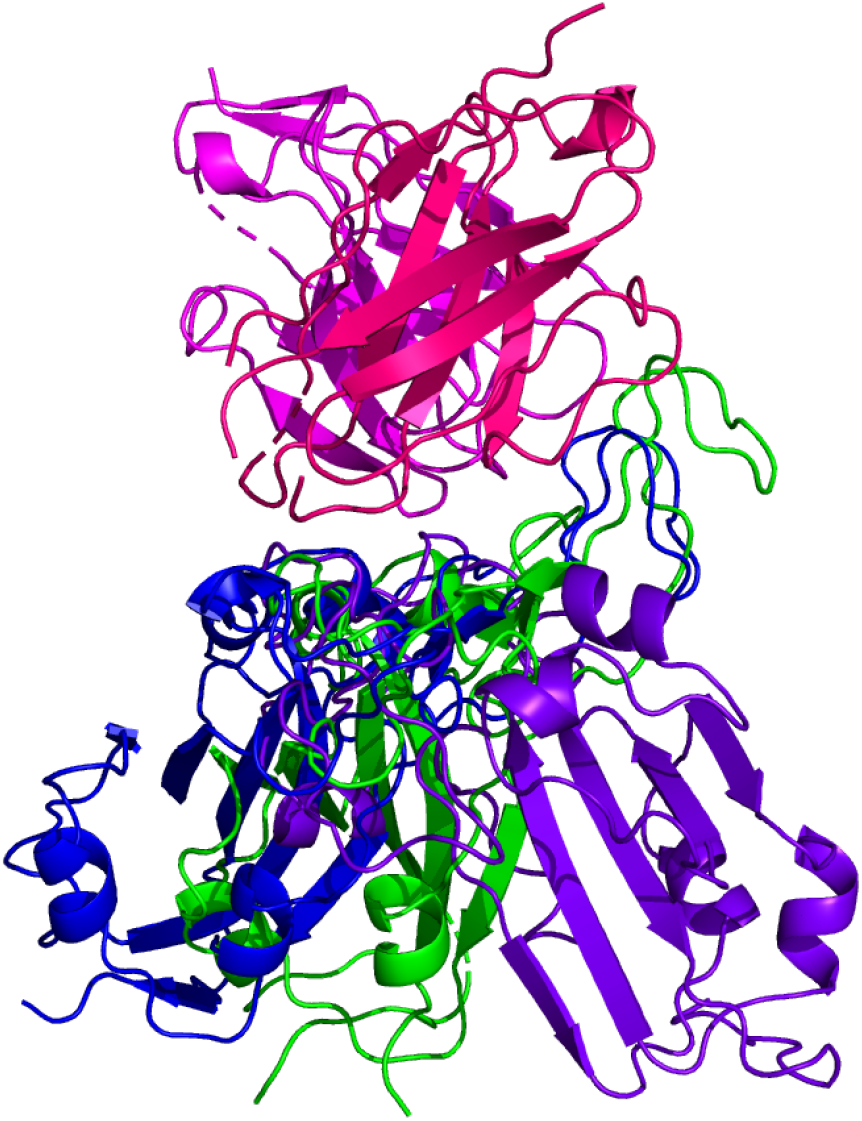
HADDOCK docking prediction using C105 (shown in magenta / pink), comparing the 6XCM RBD (shown in shown in green) vs. the Omicron (B.1.1.529) predicted RBD structures (`AlphaFold2` shown in blue, `RoseTTAFold` shown in purple).

**Fig. 8.**
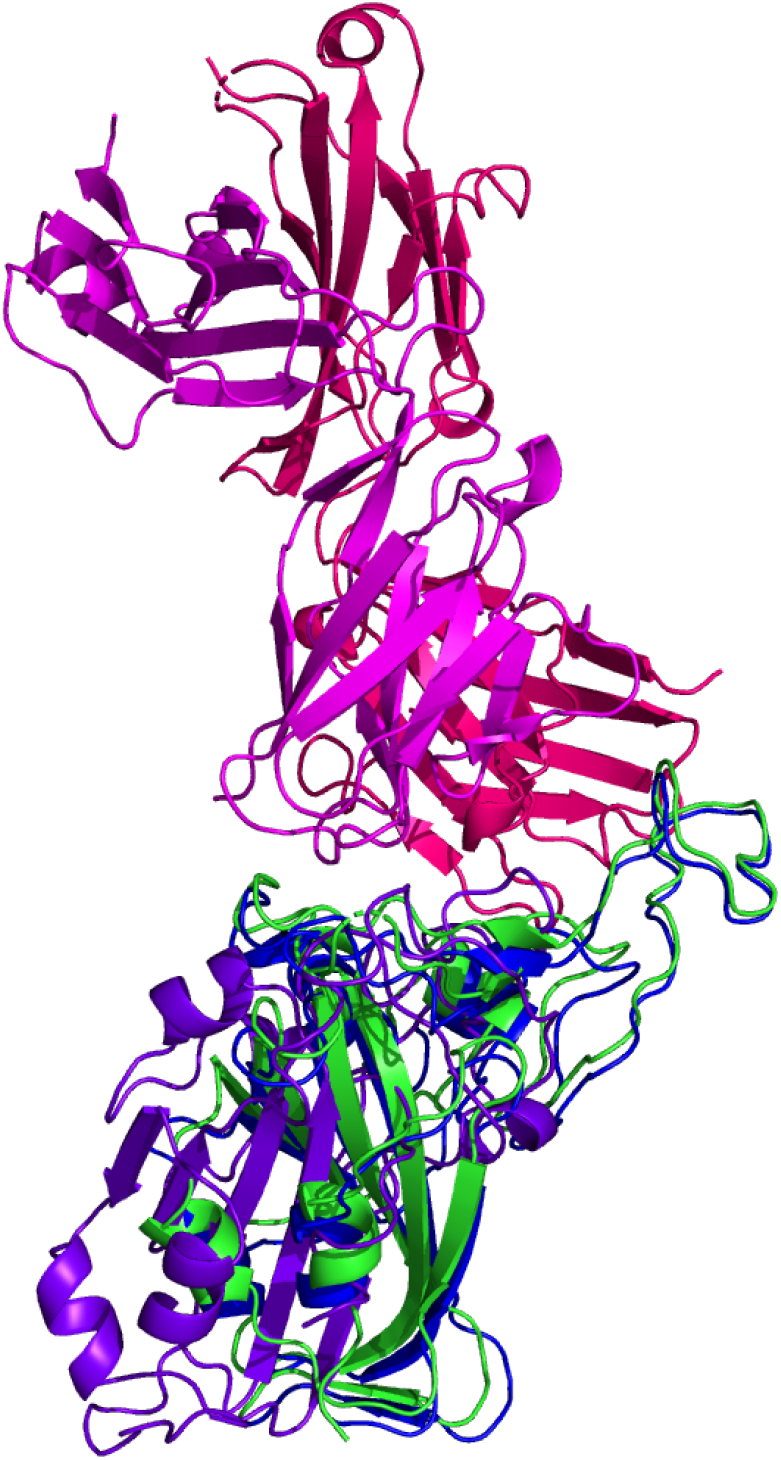
HADDOCK docking prediction using CC12.1 (shown in magenta / pink), comparing the 6XC7 RBD (shown in shown in green) vs. the Omicron (B.1.1.529) predicted RBD structures (`AlphaFold2` shown in blue, `RoseTTAFold` shown in purple).

**Fig. 9.**
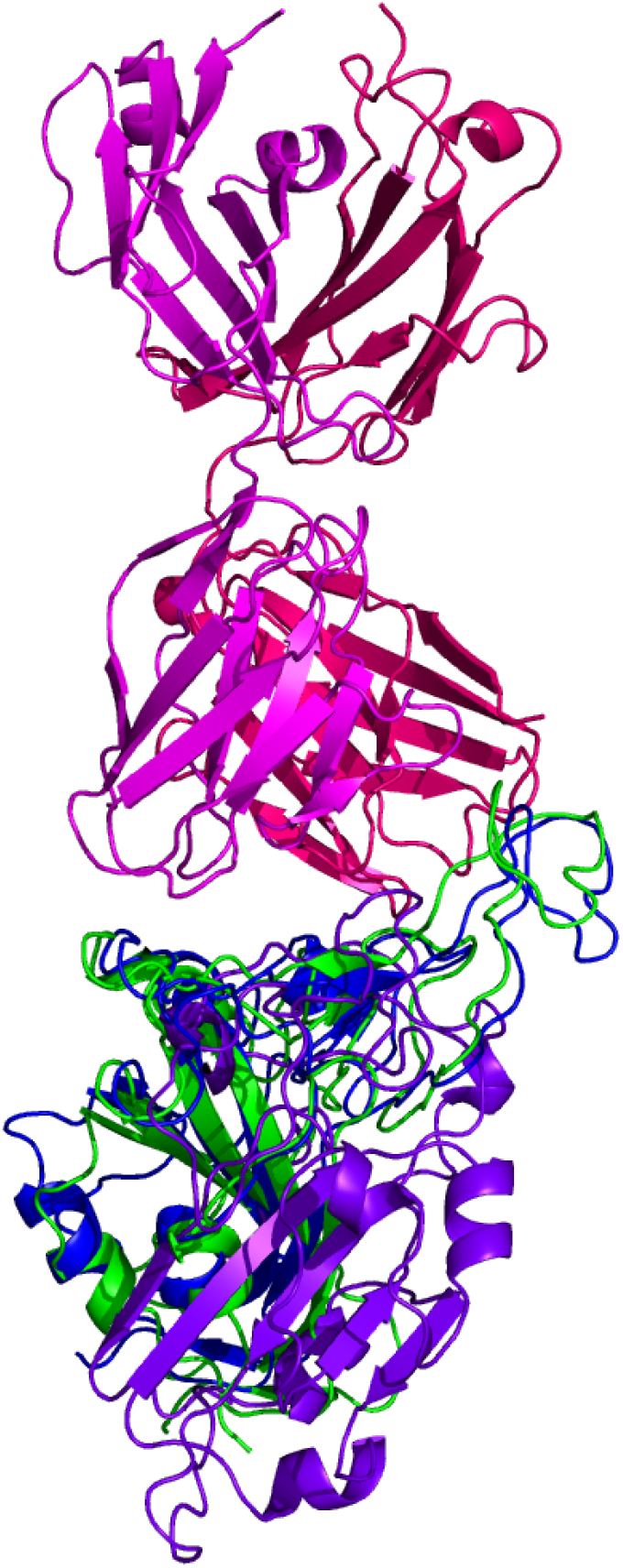
HADDOCK docking prediction using CC12.3 (shown in magenta / pink), comparing the 6XC2 RBD (shown in shown in green) vs. the Omicron (B.1.1.529) predicted RBD structures (`AlphaFold2` shown in blue, `RoseTTAFold` shown in purple).

**Fig. 10.**
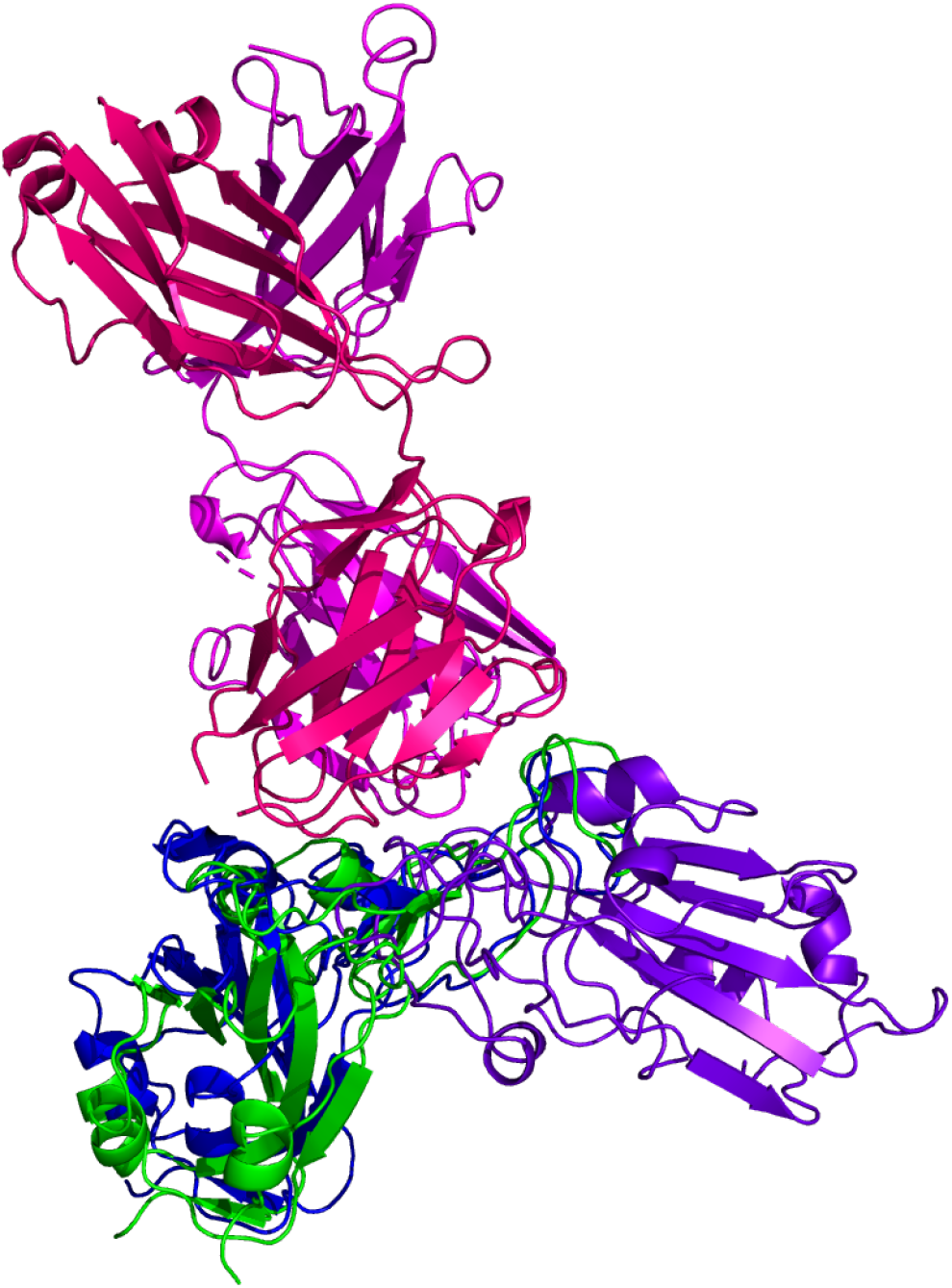
HADDOCK docking prediction using CV30 (shown in magenta / pink), comparing the 6XE1 RBD (shown in shown in green) vs. the Omicron (B.1.1.529) predicted RBD structures (`AlphaFold2` shown in blue, `RoseTTAFold` shown in purple).

**Fig. 11.**
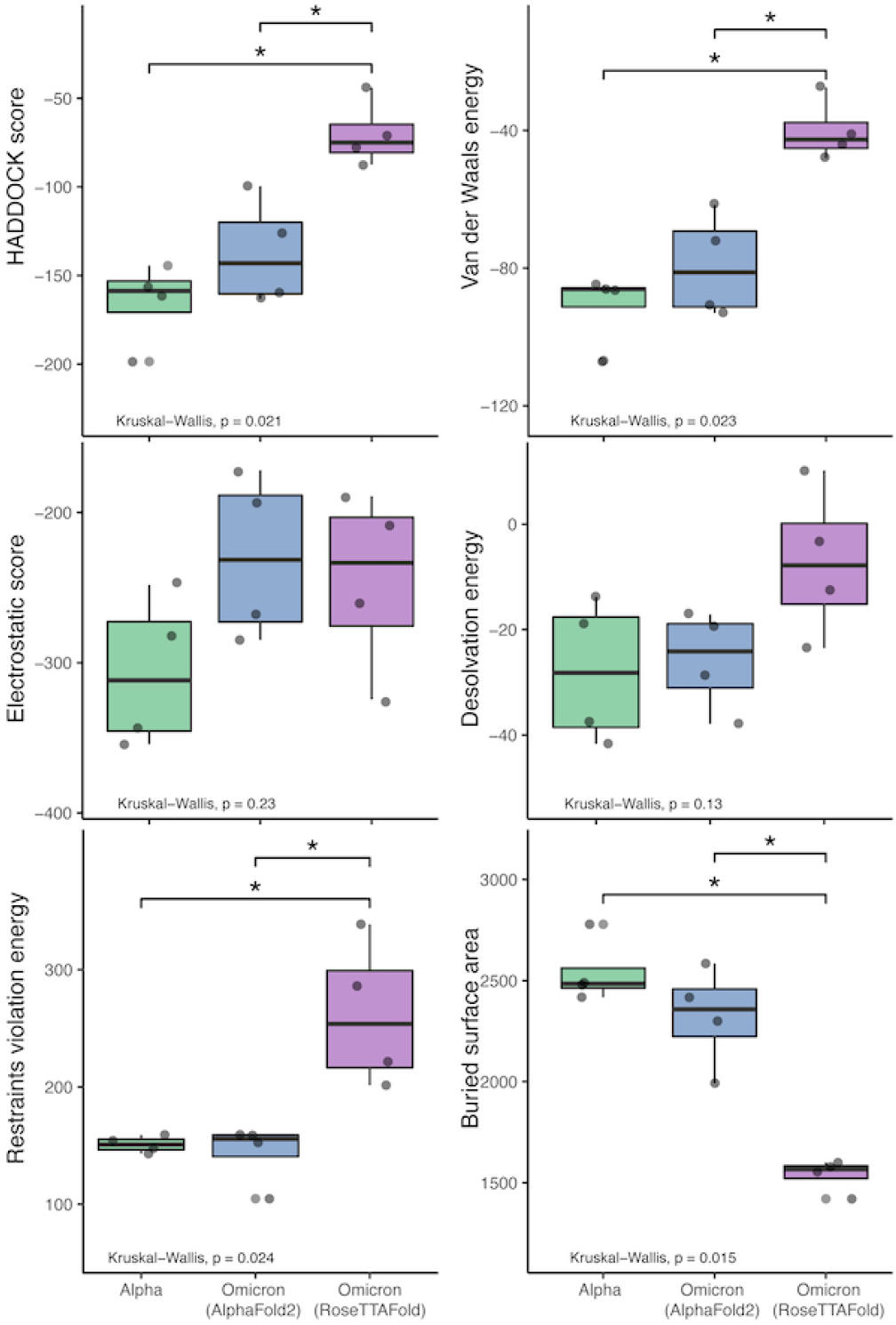
HADDOCK results comparison between the reference RBD structures and the predicted Omicron (B.1.1.529) RBD structures. The bars indicate results that are significantly different and the “*” indicates Wilcoxon Rank-Sum’s p-value *≤* 0.005).

Similarly, we see a more extreme reduction in the binding affinity of the Fab structures and the RBD structure predicted by `RoseTTAFold`. We see a drop in electrostatic interaction (increase in the electrostatic energy value) ranging from ∼ 24% to∼ 41% in three of the four interactions (with an odd ∼ 16% increase in the C105 interaction) and a consistent ∼35% to ∼43% decrease in buried surface area (increase in distance) of the RBD and the antibody Fab.

We performed Kruskal-Wallis tests with each of the `HADDOCK` metrics, which show that there is a statistically-significant difference between the three sets of binding experiments (Alpha and the two predicted Omicron sets). See Figure 11. Interestingly, performing the Wilcoxon RankSum tests on these metrics to compare the differences between the predictions and reference results shows that there is no statistically-significant difference at the *α* = 0.05 level between the `AlphaFold2` RBD structure binding and the reference structures. However, when comparing the `RoseTTAFold` RBD structure to the reference results, differences in the restraints violation energy, van der Waals energy, HADDOCK score, and buried surface area are statistically significant at the *α* = 0.05 level and the differences in desolvation energy is statistically significant at the *α* = 0.10 level. Also, the same metrics are significantly different between the `AlphaFold2` and `RoseTTAFold`, further showing that these predicted structures are quite different from one another.

#### Fab-RBD Interfacing Residues

Furthermore, when comparing residues that are interfaced between the Fab and RBD, there is agreement in that particular residues in Omicron are no longer interfacing with the antibodies analyzed in this study. In particular, residues 448N, 484A, and 494S may not interface with the Fab structure as they are in the reference RBD-Fab complexes. However, the aforementioned N501Y and S477N mutations (along with a variety of other mutations) do not appear to affect the interfacing of the residues at these positions.

This implies that there are certain positions that are more sensitive to mutations in that substitutions at these loci are more likely to affect the interface of the RBD with the antibody’s Fab (denoted by a Δ symbol in Table 7).

**Table 7.**
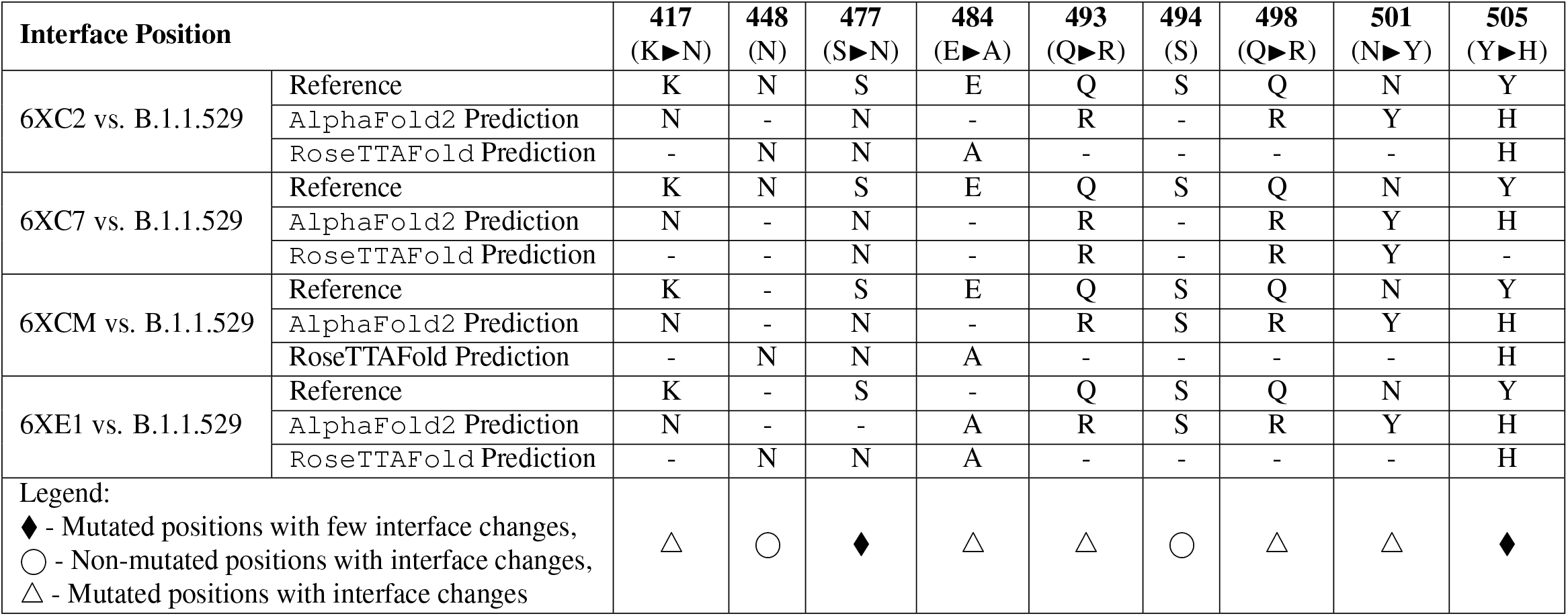
Interfacing residue changes of interest between the Fab paratope and the RBD structures. (Note that a ‘-’ means that the residue at this position no longer interfaces with the Fab structure.)

In contrast, there are other positions that have been substituted between the reference RBDs and the predicted Omicron RBDs that continue to interface with most of the Fab structures (denoted by a ♦ symbol in Table 7).

Finally, there are some residues that remain unchanged in the Omicron variant RBD structure, yet we see changes in the interfacing at these loci (denoted by a ° symbol in Table 7). This suggests that there are other mutated residues around these stable positions that may be affecting their ability to interface.

Many of these findings concur with the findings in Sharma et al. (21). Specifically, we also continue to see a reliance on residues at positions K417, S477, Q493, Q498, N501, and Y505, which are stated to increase binding affinity. Interestingly, the interfacing residues that are non-mutated N448 and S494 (denoted with a ° symbol) are rarely, if ever, listed as interfacing residues in Table 3 in Sharma et al. (21). This further supports the notion that there are residue positions that are important for interfacing with neutralizing antibodies, but that the mutations seen here in the Omicron RBD may not seriously affect this RBD-antibody interface.

### Conclusion and Discussion

While *in vitro* experiments are needed to validate these predictions, the predicted results here suggest that existing neutralizing antibodies may still bind to the mutated spike protein of the Omicron variant. However, it appears that the affinity of Omicron’s RBD for neutralizing antibodies is reduced compared to the reference RBD structures. The results of both `AlphaFold2` and `RoseTTAFold` suggest that antibodies elicited from previous infection will provide at least some protection against Omicron. Additionally, these results indicate that the SARS-CoV-2 Omicron variant will note completely evade vaccines based on the spike protein.

Though there are many mutations in the RBD of Omicron, the predicted structure from `AlphaFold2` suggests these mutations do not appear to be causing any large conformational change that would totally evade antibody interaction. However, we do see some amino acid substitutions to different, longer side chained residues at the binding site. This result may be due to the slightly more distant interaction with the antibody and therefore may reduce the binding affinity. The results of `RoseTTAFold` suggest that the mutations have a different effect on the overall 3D structure of the Omicron RBD. The conformational change seen in this structure may contribute to antibody evasion or more severely reduce antibody binding affinity.

`AlphaFold2` has not been validated for predicting the effect of mutations and is not expected to produce a completely unfolded structure if the Omicron sequence contains any destabilising point mutations. This may explain the extremely similar backbone structure to the reference RBD structures.

Given that the antibody docking results between `AlphaFold2` and `RoseTTAFold` are statistically different, this further posits that this potential conformational change seen in the `RoseTTAFold` RBD may directly affect the binding of neutralizing antibodies.

Though the predicted RBD structures between `AlphaFold2` and `RoseTTAFold` are quite different, the `HADDOCK` antibody docking results seem to converge on specific positions with which the Fab will interface. S477N and Y505H may be useful positions on which to focus for future vaccine design or mutational surveillance in new variants.

Further analyses are needed using a broader range of different classes of antibodies, including therapeutic antibodies and neutralizing antibodies that bind to other locations on the spike protein. Recent preprint articles show more drastic reductions in the binding affinity of some other antibodies like CB6, a neutralizing antibody similar to ones used in this study, as well as a variety of therapeutic antibodies (22, 23). Thus, while we fail to see complete evasion of the antibodies used in this study, there are far more Omicron-antibody interactions to be understood.

Once a true structure of the Omicron RBD is determined, it will be of interest to compare the true structure to both of the predicted RBD structures from `AlphaFold2` and `RoseTTAFold`, mainly to see if a large conformational change occurs (as is seen in the `RoseTTAFold`) or if the overall backbone/3D structure is very similar to the reference structures used in this study (as is seen in the `AlphaFold2`). In addition, it will be necessary to validate the antibody interactions predicted using `HADDOCK` with true, experimentallyderived binding measurements.

Determining the actual structure of a protein is a time-consuming process. Further, quantifying protein-protein interactions (like spike-to-antibody interactions) are also experimentally difficult to perform *in vitro*. Given the public health urgency in understanding the impacts of new SARS-CoV-2 variants quickly requires that we act quicker than is possible in a lab. Thus, *in silico* predictive tools like `AlphaFold2`, `RoseTTAFold`, and `HADDOCK` are important for quickly understanding the biochemistry of variants and can help us to infer the epidemiological implications of the variant.

## Supplementary Materials

All data, scripts, and results from this work are available at GitHub.com/colbyford/SARS-CoV-2_B.1.1.529_SpikeRBD_Predictions (24).

## ACKNOWLEDGEMENTS

We acknowledge these entities at UNC Charlotte: The Department of Bioinformatics and Genomics, The Bioinformatics Research Center, the College of Computing and Informatics, The College of Liberal Arts and Sciences, Research and Economic Development, University Research Computing, and the Graduate School. We also thank the Belk family for support.

Also, we appreciate the initial feedback from members of the Google DeepMind team on the usage of `AlphaFold2` for point mutations. This prompted our further investigation and comparison with `RoseTTAFold` and the differing results we now show in this study.

## Footnotes

1 *InterafaceResidues* Function Documentation: https://pymolwiki.org/index.php/InterfaceResidues

